# The neural G protein Gα_o_ tagged with GFP at an internal loop is functional in *C. elegans*

**DOI:** 10.1101/2021.02.16.431479

**Authors:** Santosh Kumar, Andrew C. Olson, Michael R. Koelle

**Author notes:** Present address: Department of Biotechnology, Panjab University, Chandigarh 160014, India (Santosh Kumar). These authors contributed equally to this work (Santosh Kumar, Andrew C. Olson). Co-corresponding authors: 300 Cedar Street SHM CE30, Dept. of MB&B, Yale University Medical School, New Haven, CT 06520,., Department of Biotechnology, Panjab University, BMS Block I, Sector 25, Chandigarh 160014, India.

## Abstract

Gα_o_ is the alpha subunit of the major heterotrimeric G protein in neurons and mediates signaling by every known neurotransmitter, yet the signaling mechanisms activated by Gα_o_ remain to be fully elucidated. Genetic analysis in *Caenorhabditis elegans* has shown that Gα_o_ signaling inhibits neuronal activity and neurotransmitter release, but studies of the molecular mechanisms underlying these effects have been limited by lack of tools to complement genetic studies with other experimental approaches. Here we demonstrate that inserting the green fluorescent protein (GFP) into an internal loop of the Gα_o_ protein results in a tagged protein that is functional *in vivo* and that facilitates cell biological and biochemical studies of Gα_o_. Transgenic expression of Gα_o_-GFP rescues the defects caused by loss of endogenous Gα_o_ in assays of egg laying and locomotion behaviors. Defects in body morphology caused by loss of Gα_o_ are also rescued by Gα_o_-GFP. The Gα_o_-GFP protein is localized to the plasma membrane of neurons, mimicking localization of endogenous Gα_o_. Using GFP as an epitope tag, Gα_o_-GFP can be immunoprecipitated from *C. elegans* lysates to purify Gα_o_ protein complexes. The Gα_o_-GFP transgene reported in this study enables studies involving *in vivo* localization and biochemical purification of Gα_o_ to complement the already well-developed genetic analysis of Gα_o_ signaling.

## Introduction

Gα_o_, the α subunit of the most abundant heterotrimeric G protein in the brain (Sternweiss and Robishaw, 1984), is in every neuron and can be activated by G protein coupled receptors for every neurotransmitter tested (Jiang et al., 2001). *C. elegans* has a Gα_o_ ortholog named GOA-1 that is >80% identical to mammalian Gα_o_ and that is expressed in most or all neurons. GOA-1 has been shown by genetic analysis to inhibit neurotransmitter release and/or neural activity (Mendel et al., 1995; Ségalat et al., 1995; Nurrish et al. 1999, Ravi et al., 2020), but the molecular mechanisms by which Gα_o_ signals to have these effects remain to be fully defined. While activated Gα_o_ releases Gβγ subunits to regulate specific potassium and calcium channels (Lüscher and Slesinger, 2010; Proft and Weiss, 2015), genetic studies in *C. elegans* suggest that signaling through Gβγ is not likely the sole mechanism by which Gα_o_ has its physiological effects (Koelle, 2018). It remains unclear if activated Gα_o,_ like all other Gα proteins in animal cells, may itself binds target “effector” proteins to propagate a signal.

A method to fuse Gα_o_ to fluorescent proteins and/or epitope tags without disrupting its function would enable new experimental approaches to help resolve unanswered questions about Gα_o_ signaling. For example, Gα_o_-GFP fusion proteins could be visualized in real time in living cells for cell biological studies, and anti-GFP antibodies could be used to immunopurify Gα_o_ protein complexes for biochemical analysis. The challenge to this approach is that tags at the N- or C-termini would likely disrupt Gα_o_ function since Gα proteins use their N- and C-termini to interact with receptors, Gβγ subunits, and membranes (Hynes et al., 2004). Thus, efforts to functionally tag Gα proteins have focused on inserting fluorescent proteins at internal sites. Internally-tagged Gα proteins have been shown to be activated by G protein coupled receptors when co-overexpressed in cultured cells with both a receptor and Gβγ subunits (Hughes et al., 2001; Yu and Rasenick, 2002; Bünemann et al., 2003; Galés et al., 2005; Lazar et al., 2011); however, some internal insertions alter Gα function, and overexpressed receptors can promiscuously activate Gα proteins they would not otherwise activate (Gibson and Gilman, 2006). Still, some tagged Gα proteins may be fully functional: in the yeast *Saccharomyces cerevisiae* and in the slime mold *Dictyostelium discoideum*, an internally-tagged Gα protein can replace the untagged Gα to support physiological functions that depend on activation by a single endogenous receptor (Janetopoulos et al., 2001; Yi et al., 2003). A remaining question is whether a tagged Gα protein could be fully functional in a metazoan, where it must mediate signaling from many different receptors to control diverse, tissue-specific physiological functions.

Here, we demonstrate that *C. elegans* Gα_o_ with GFP inserted into an internal loop, when expressed at normal levels in the animal, rescues multiple defects in behavior and development caused by loss of native Gα_o_. We show that this tagged protein can be used to visualize Gα_o_ subcellular localization in living animals and to purify both inactive and activated Gα_o_ protein complexes from *C. elegans* lysates.

## Materials and Methods

All reagents were from Sigma-Aldrich (St. Louis, MO) unless otherwise indicated.

### Strains and culture

*C. elegans* strains were cultured at 20°C on NGM agar plates with *E. coli* strain OP50 as a nutrition source (Brenner 1974). All strains were derived from the wild-type strain N2. Generation of transgenic animals and genetic crosses were by standard methods (Evans 2006; Fay 2013). Table 1 shows a list of *C. elegans* strains used in this study.

**Table 1.** *C. elegans* strains used in this study.

| Strain | Genotype | Feature | Source |
| --- | --- | --- | --- |
| N2 | Wild type |  | Brenner, 1974 |
| LX1691 | *unc-119(ed3) III* | Recipient strain for miniMos transgenes. Made by outcrossing HT1593 (from the *Caenorhabditis* Genetics Center) 4X to N2. | This study |
| EG6699 | *ttTi5605 II; unc-119(ed9) III; oxEx1578* | Unc progeny (which have lost the *unc-119*-rescuing transgene *oxEx1578*) are recipients for single-copy transgene insertion into the *ttTi5606* Mos1 locus. | *Caenorhabditis* Genetics Center |
| LX2060 | *vsSi32 III unc-119(ed3) III* | *goa-1::gfp* strain. | This study |
| LX2404 | *vsSi39 II* | *goa-1(Q205L)::gfp* strain. | This study |
| JT734 | *goa-1 (sa734) I* | *goa-1* null mutant | Robatzek and Thomas, 2000 |
| LX2071 | *goa-1(sa734) I; vsSi32 III* | *goa-1::gfp* in *goa-1* null background | This study |

### *goa-1::gfp* plasmid construction

A plasmid to express internally GFP-tagged GOA-1 in *C. elegans* was generated by first inserting a 9.0 kb *C. elegans* genomic fragment containing the *goa-1* gene into a pBluescript vector and engineering in an SpeI restriction site between the *goa-1* codons for T^117^ and E^118^. The GFP coding region containing artificial introns was PCR-amplified from the vector pPD95.69 (Addgene plasmid #1491) using primers to add segments encoding SGGGGS and SGGGTS to flank the N- and C-termini of GFP, respectively, and the resulting *gfp* cassette was inserted into the SpeI site of the *goa-1* clone. The resulting plasmid was named pMK376. To generate a clone suitable for miniMos single-copy insertion into the *C. elegans* genome (Frøkjær-Jensen et al., 2014), we amplified the GOA-1-GFP coding region from pMK376 along with 4987 bp of 5’ promoter and 432 bp of 3’ UTR using primers mini909FWD (5’-gagatt**ctgcag**gaattccaactgaatttagatttttaaagt-3’) and mini909REV (5’-agatt**aggcct**ggagtcttttcacccatacttccggaataa-3’), which added StuI and PstI restriction sites (boldface) at the 5’ and 3’ ends of the amplicon, respectively. The amplicon was then digested and ligated into the pCFJ909 miniMos vector (Frøkjær-Jensen et al., 2014) using the StuI and PstI restriction sites. The resulting plasmid was named pAO8.

The Q205L mutation was engineered into *goa-1::gfp* gene pMK376 using the GeneArt Site-Directed Mutagenesis PLUS Kit (Invitrogen) and the primers SKB48 (5’-tgtgggaggtctgagatcagaaag-3’) and SKB49 (5’-tcgaacaatctgtaaatattc-3’). To generate a clone for site-specific single-copy insertion into the *C. elegans* genome, the *goa-1(Q205L)::gfp* cassette was then amplified using primer pair SKB46 (5’-agatacctaggggagtcttttcacccatacttccggaataa-3’) and SKB47 (5’-ctcacttaaggaattccaactgaatttagatttttaaagt-3’) to add AvrII and AflII restriction sites, and this fragment was subsequently subcloned into the MosSCI insertion vector pCFJ151 (Frøkjær-Jensen et al., 2008) digested with AvrII and AflII. The resulting plasmid was named pSKB21.

### Single-copy *goa-1::gfp* transgenic strains

The *goa-1::gfp* single-copy miniMos transgene *vsSi32* was generated as described by Frøkjær-Jensen et al. (2014) by injecting into LX1691 *unc-119 (ed3) III* animals the *goa-1::gfp* plasmid pAO8 at 15 ng/μl, with the Mos1 transposase plasmid pCFJ601 at 50 ng/μl, and marker plasmids pCFJ90 at 2.5 ng/μl, pCFJ104 at 10 ng/μl, pGH8 at 10 ng/μl, and pMA122 at 10 ng/μl. Inverse PCR was used to determine the transgene integrated in the left arm of chromosome III between the sequences 5’-tttactgcatactgaacaacaggggaaaagggg-3’ and 5’-tagaattagctgtaagacggcgtctaggttttgca-3’. The *goa-1(Q205L)::gfp* MosSCI single-copy transgene *vsSi39* was inserted at the *ttTi5605* Mos1 locus on chromosome II as described by Frøkjær-Jensen et al. (2008) by injecting into Unc progeny of EG6699 *ttTi5605 II; unc-119(ed9) III; oxEx1578* animals a mix of the *goa-1(Q205L)::gfp* plasmid pSKB21 at 50ng/ul, the Mos1 transposase plasmid pCFJ601 at 50 ng/μl, and marker plasmids pCFJ90 at 2.5 ng/μl, pCFJ104 at 5 ng/μl, pGH8 at 10 ng/μl, and pMA122 at 29 ng/μl. The initially generated transgenic animals were outcrossed four times to wild-type N2 animals to remove background mutati0ns. The resulting strain was named LX2404.

### Confocal imaging

Worms were on mounted on 2% agarose pads containing 120 mm Optiprep (Sigma Millipore) to reduce refractive index mismatch (Boothe et al., 2017) on premium microscope slides Superfrost (Thermo Fisher Scientific), and a 22 × 22–1 microscope cover glass (Fisher Scientific) was placed on top of the agarose pad. Worms were anesthetized using a drop of 150 mm sodium azide (Sigma Millipore) with 120 mm Optiprep. Z-stack confocal images of 24 hour old larvae were taken on a Zeiss LSM 880 microscope using a 40X objective lens.

### GOA-1 antibody

The affinity-purified rabbit anti-GOA-1 polyclonal antibody used was from Chase et al. (2001). Whole mount stains of *C. elegans* were performed as described by Finney and Ruvkun (1990).

### Behavioral and worm length assays

Quantitation of unlaid eggs and staging of laid eggs were performed using 30 hour post-L4 adult animals as described in Chase and Koelle (2004). For analysis of worm tracks, reversal-touch behavior, and worm length, worms were staged 24 hours post-L4 and transferred to an NGM agar plate with a thin lawn of OP50 bacteria for imaging. Imaging began 2-20 minutes after transfer of the worm to the new plate and was carried out using a Leica M165FC microscope equipped with a digital camera. For measurements of worm length, >10 second digital video recordings of worms were analyzed using WormLab software from MBF Bioscience. Reversal-touch behavior was defined as a reversal during which the worms bends deeply enough that it contacts itself. Usually the tail or head touches the body, but in some cases two sections of the midbody can touch each other during very deep bends. Reversal-touch behavior was scored by placing a single worm staged 24 hours post-L4 on a new NGM plate with a lawn of OP50 bacteria, waiting 1-10 minutes, and then counting the number of reversal-touches made by the worm during a 15-second time period.

### Statistical analysis

Error bars shown in the graphs in Figures 2–3 represent 95% confidence intervals. All statistical analyses were done using GraphPad Prism version 9.0.1 software. The early-stage egg assay data set was analyzed using Fisher’s exact test with two-sided P-values. The remaining data sets were analyzed using one-way ANOVA with Šídák’s multiple comparisons test.

**Figure 1.**
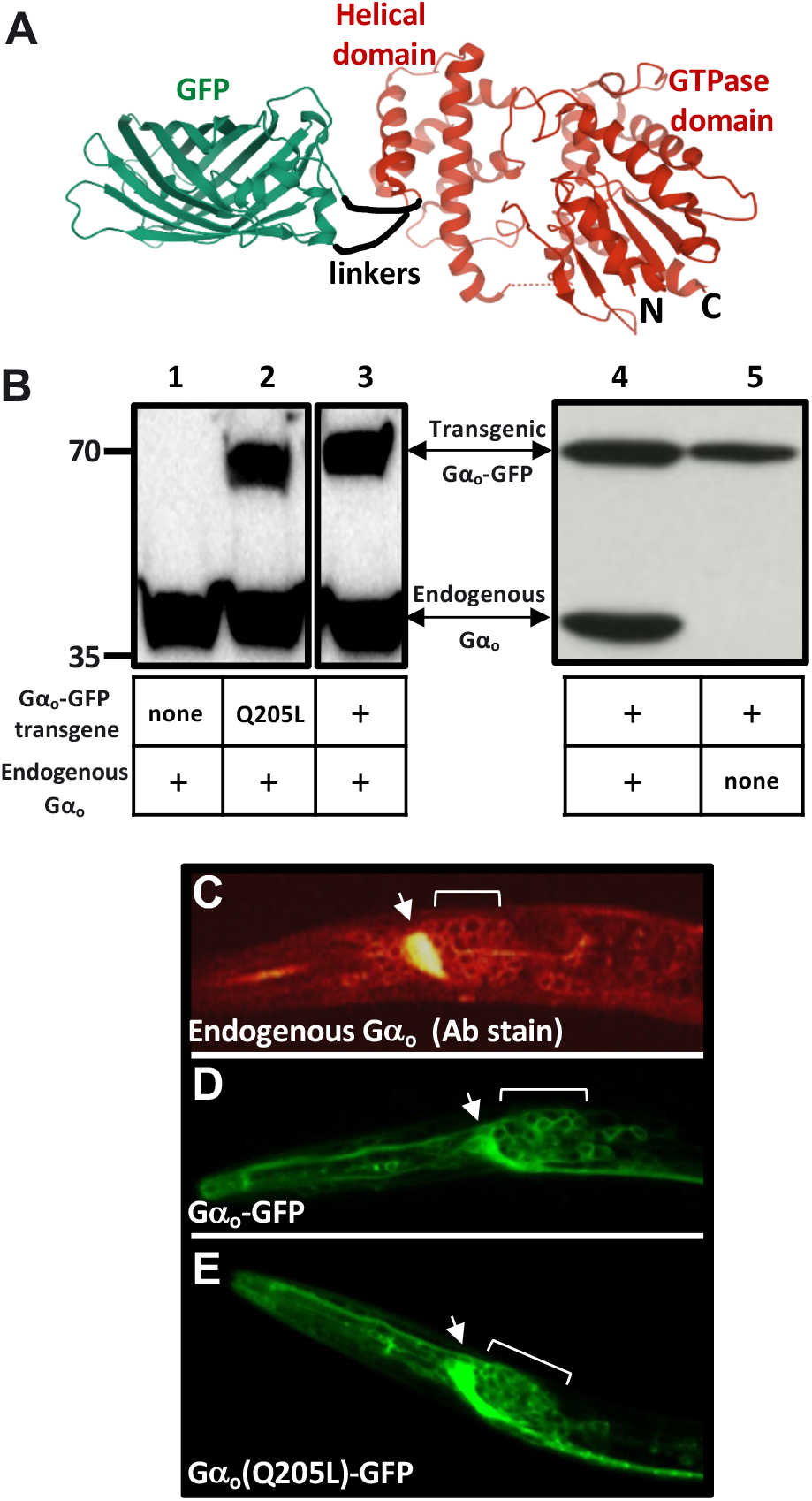
Design and expression of Gα_o_-GFP in *C. elegans*. (A) Design of Gα_o_-GFP. Structures of mammalian Gα_o_ (red, PDB entry 3c7k) and GFP (green, PDB entry 1GFL) were rendered with Mol* (Sehnal et al 2018). The designed fusion protein has six amino acid linkers (black) connecting the GFP N- and C-termini to amino acids T117 and E118 of *C. elegans* Gα_o_. (B) Western blots of *C. elegans* protein lysates probed with an antibody to Gα_o_ (Patikoglou and Koelle 2002) that detects both endogenous Gα_o_ and transgenically-expressed Gα_o_-GFP. Lanes 1-3 are from the same gel, while lanes 4-5 are from a separate gel. Positions of molecular weight markers (in kDa) ran at positions shown at left. Protein lysates were from five *C. elegans* strains in which Gα_o_-GFP transgenes and a null mutation in the endogenous Gα_o_ gene were present in the combinations indicated. The strain analyzed in lane 2 expressed Gα_o_-GFP carrying a Q205L mutation that disrupts Gα_o_ GTPase activity and thus renders the G protein constitutively active. (C-E) Localization of endogenous Gα_o_ protein by anti-Gα_o_ antibody stain (C) compared to localization of Gα_o_-GFP (D) and Gα_o_(Q205L)-GFP (E) by GFP fluorescence. Shown are confocal images of optical sections through the heads of *C. elegans* larvae, with anterior left and ventral down. Arrows point to the nerve ring, a bundle of neural processes containing many synapses; brackets indicate the neural ganglia posterior to the nerve ring, containing the cell bodies of many neurons. Rings of fluorescence in the neural ganglia suggest that Gα_o_ is localized on the plasma membrane of neural cell bodies. Antibody staining is absent in Gα_o_ null mutant animals stained in parallel with the same Gα_o_ antibody, demonstrating that the stain is specific for Gα_o_ (Gotta and Ahringer 2001, and data not shown).

**Figure 2.**
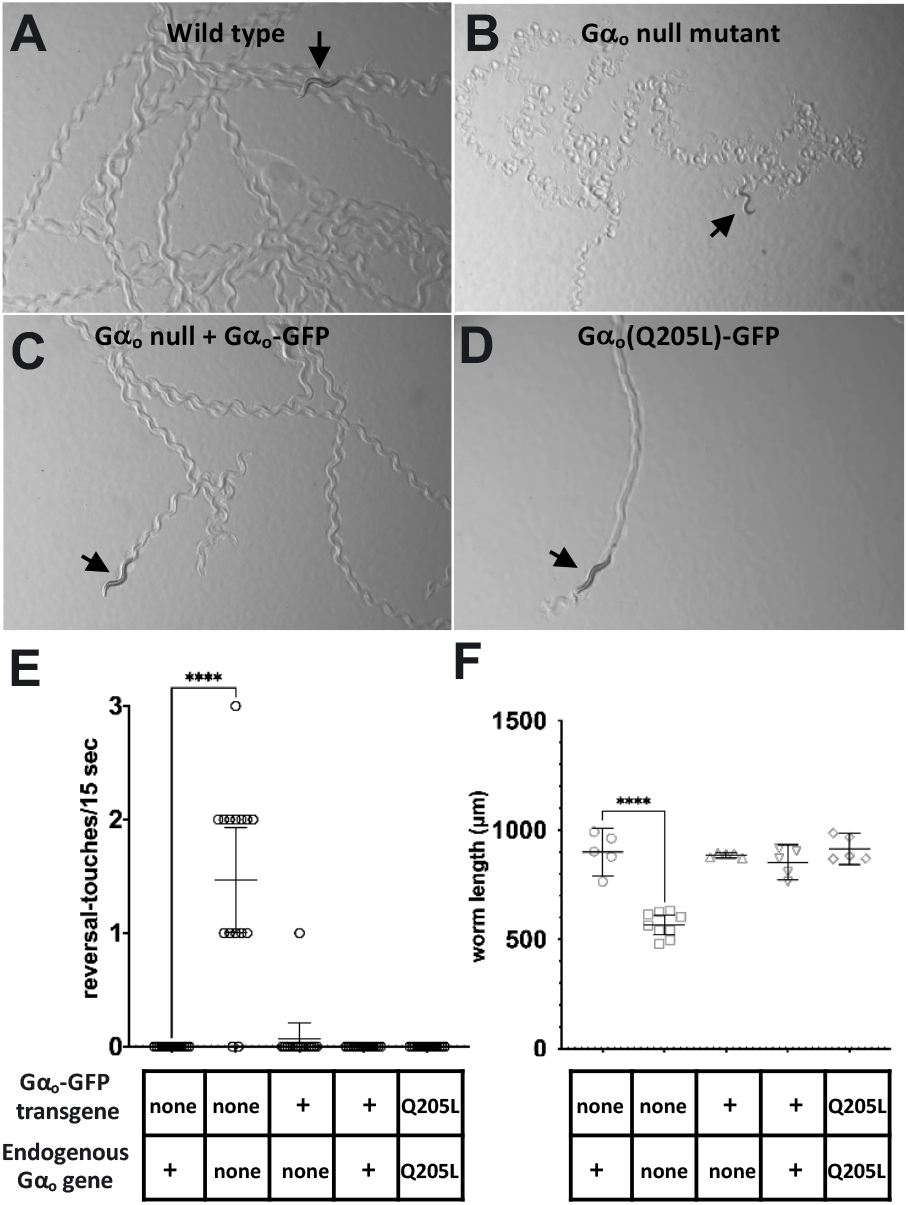
Gα_o_-GFP rescues the locomotion and body morphology defects of a Gα_o_ null mutant. (A-D) Photographs of single worms of the indicated genotypes showing tracks left by locomotion over an agar surface. Arrows indicate current positions of the worms. Sinusoidal tracks left by the wild-type worm (A) show the pattern of normal locomotion, whereas the Gα_o_ null worm in (B) is smaller and has left tracks that reflect abnormal locomotion. Expression of Gα_o_-GFP in the Gα_o_ null background restores locomotion to a more normal pattern (C), while expression of activated Gao(Q205L)-GFP in a wild-type background (D) induces a gain-of-function phenotype evidenced by a track that reflects abnormally shallow body bends. (E) Animals of the indicated genotypes (n=15 per genotype) were video recorded and reviewed to quantitate the frequency of episodes of backwards locomotion that included a body bend so deep that the animal touched its own body. (F) Videos of adult worms of the same genotypes (n≥5 per genotype) were analyzed with WormLab software to measure their body length. **** indicates p<0.0001; error bars represent 95% confidence intervals.

**Figure 3.**
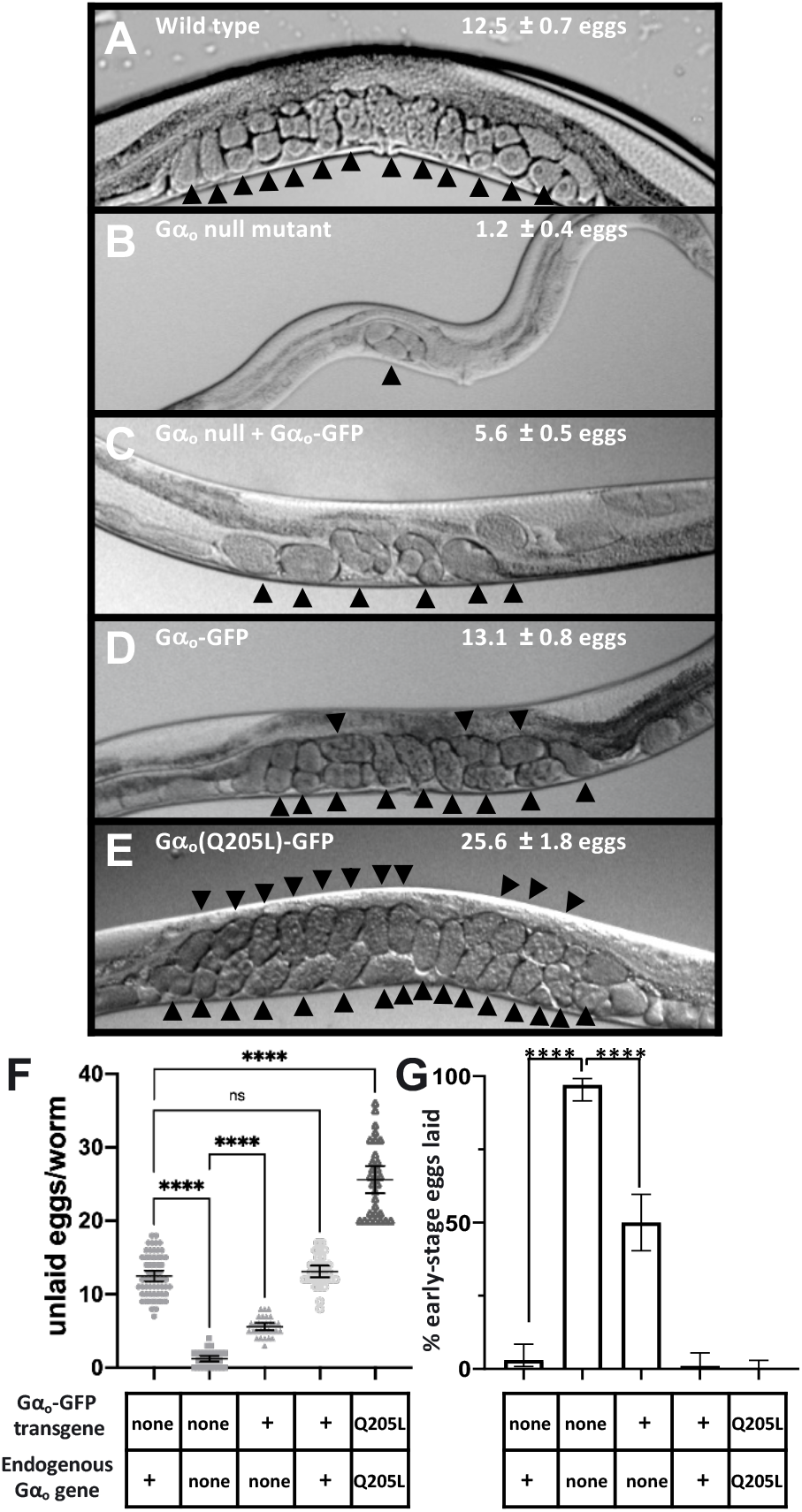
Gα_o_-GFP partially rescues egg-laying behavior defects of a Gα_o_ null mutant. (A-E) Photographs of worms of the indicated genotypes showing unlaid eggs within the mid-body region. Each unlaid egg is indicated by an arrowhead, and the average number of unlaid eggs and 95% confidence intervals for each genotype are shown on the corresponding photographs. The scrawny body morphology of a Gα_o_ null mutant (B) compared to that of a wild-type animal (A) can also be seen. (C) Quantitation of unlaid eggs, n≥30 for each strain. (D) Percent of freshly laid eggs at early stages of development (≤8 cells), a measure of hyperactive egg-laying behavior. ****, p<0.0001; ns, not significant (p>0.05); error bars represent 95% confidence intervals.

### Immunoprecipitation of GOA-1::GFP

Worm lysates were prepared as described previously (Porter and Koelle, 2010) with some modifications. Briefly, *C. elegans* were grown in 20 ml liquid cultures at 20° C and worms were isolated by flotation on 30% sucrose. Packed worm pellets (~500-600 μl) were resuspended in 4 ml lysis buffer (50 mM HEPES pH7.4, 100 mM NaCl, 1 mM EDTA, 3 mM EGTA, 10 mM MgCl_2_, 1 mM DTT, 1% Triton X-100 and complete protease inhibitor cocktail (Roche #04693159001). Resuspended worm pellets were homogenized by passing them two times through a French press (Spectronic Instruments, model number FA078) at 13000 PSI. The resulting lysates were centrifuged at 100,000Xg for 30-60 min at 4°C in an Optima TLX tabletop ultracentrifuge using a TLA-110 rotor (Beckman Coulter, Fullerton, CA). The clarified supernatants were removed and transferred into the new tubes. The protein concentrations were determined by Bio-Rad protein assay. Whole worm lysates were aliquoted and snap frozen in liquid nitrogen for storage at −80°C until use.

Immunoprecipitation was performed using Pierce crosslink immunoprecipitation kits (Pierce #26147) and the buffers therein, per manufacturer’s instructions with some modifications. All spins in this procedure to separate supernatants from beads were at 3,000 RPM for 1 minute in a microcentrifuge equipped with a swinging bucket rotor. To prepare beads sufficient for immunoprecipitating three samples, 90 μl of 50% protein A/G agarose slurry/sample was washed three times with1 ml phosphate buffered saline (PBS, 10 mM sodium phosphate, 0.15 M NaCl, pH 7.5 and then incubated at room temperature for 2 hours on a rotary mixer at room temperature with 10 μg mouse monoclonal anti-GFP antibody (Rockland #600-301-215) diluted in 500 μl PBS. Beads were washed three times with PBS followed by incubation with Pierce DSS crosslinker working solution (150 μl) at room temperature for 1 hour on a rotary mixer. The supernatant was discarded and the beads were washed twice with Pierce elution buffer followed by three washes with IP buffer (50 mM HEPES pH 7.4, 100 mM NaCl, 1 mM EDTA, 3 mM EGTA, 10 mM MgCl_2_ and 1% Triton X-100). While the anti-GFP beads were being prepared, protein lysates were pre-cleared: for each immunoprecipitation sample, 1 ml of 4 mg/ml worm protein lysate was incubated with 15 μl packed protein A/G beads (prewashed 3X in PBS) at 4°C for 1 hour on a rotary mixer. Then for each immunoprecipitation, 1 ml of 4 mg/ml protein pre-cleared protein lysate was incubated with 15 μl packed anti-GFP antibody cross-linked beads at 4° C for 2 hours on a rotary mixer. For in vitro activation (Figure 4E), at the beginning of this 2 hour incubation, 100 μM GDP or GTPγS was added to the protein lysate. After removing the supernatant, the beads bearing the immunoprecipitated products were washed four times with IP buffer at 4° C and eluted with 50μl of 2X LDS loading buffer (Invitrogen #NP0007) at 55° C for 30 min. The supernatants were transferred into the new tubes and 5% (by volume) β-mercaptoethanol was added and the tubes were incubated at 55° C for 15 min. The supernatants were collected, and samples were processed for SDS-PAGE gel electrophoresis and western blotting.

**Figure 4.**
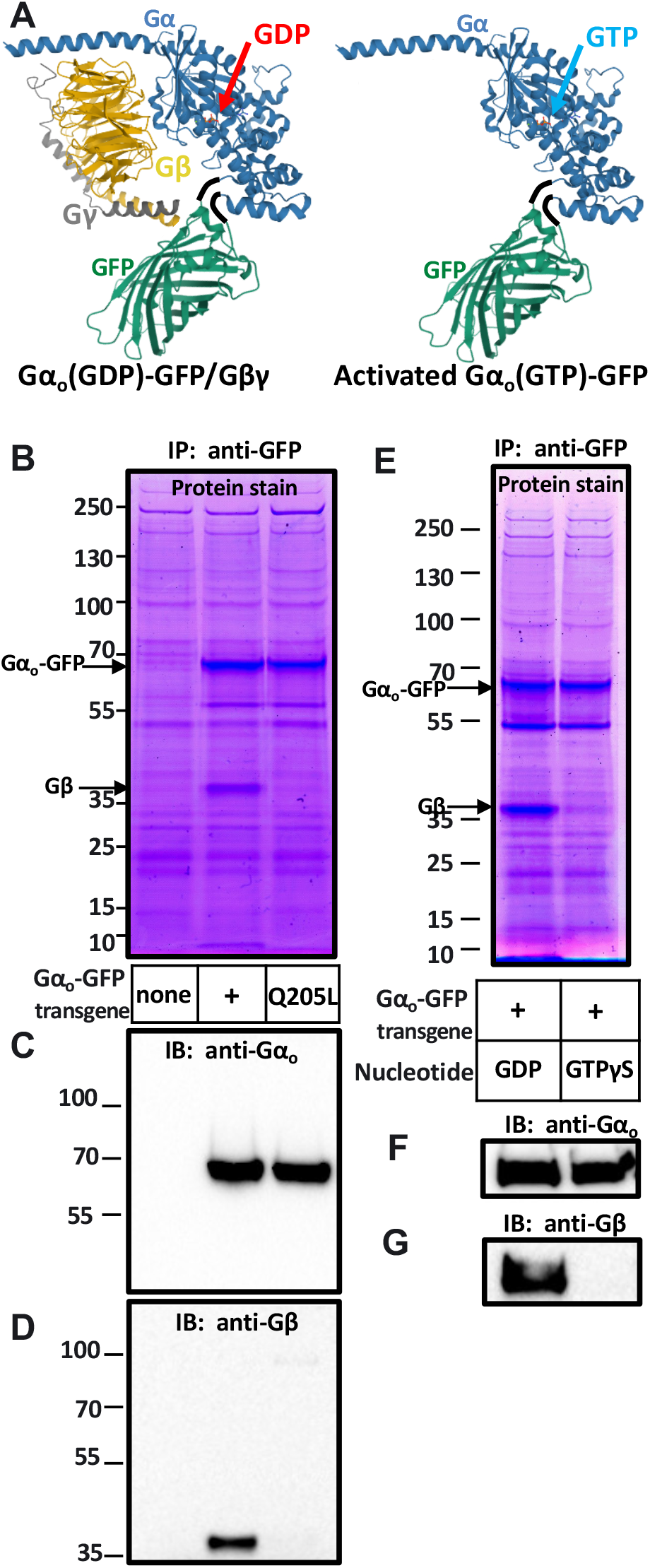
Immunoprecipitation of inactive and activated Gα_o_-GFP protein complexes from worm protein lysates. (A) Models of inactive (left) and active Gα_o_-GFP. Models were rendered with Mol* (Sehnal et al 2018) and based on mammalian Gα_s_-GDP/βγ 35(PDB entry 6EG8) with the alpha subunit (blue) connected by flexible six-amino acid linkers (black) to GFP (green, PDB entry 1GFL). Inactive Gα-GFP-GDP (left) is in a complex with Gβγ subunits, while active Gα-GFP-GTP (right) is not. (B-D) Immunoprecipitation of inactive and in *vivo*-activated Gα_o_. Proteins were immunoprecipitated with an anti-GFP antibody from whole-worm lysates of strains carrying no transgene (left lane, a control to show non-specific background proteins), the Gα_o_-GFP transgene (center lane), or the activated Gα_o_(Q205L)-GFP transgene (right lane). Proteins were resolved by gel electrophoresis, with the mobility of molecular weight marker (in kDa) shown at left. (B) Precipitated proteins were visualized using Imperial protein stain. Gα_o_-GFP and Gβ appeared as most prominent proteins precipitated, with Gβ absent from the activated Gα complex (right lane). Gγ is a small protein expected to run at the dye-front in the center lane. The identities of the prominent bands in (B) are confirmed by Western blots of the same protein samples probed with anti-Gα_o_ (C) and anti-Gβ (D) antibodies. (E-G) Immunoprecipitation of inactive and in *vitro*-activated Gα_o_. Whole worm protein lysate from the strain carrying Gα_o_-GFP (same lysate used for the middle lanes in (B-D)) was treated with GDP to preserve the inactive state of Gα_o_-GFP or with the non-hydrolysable GTP analog GTPγS to activate Gα_o_-GFP. Proteins were separated by gel electrophoresis and visualized with Imperial protein stain (B), or by Western blotting with anti-Gα_o_ (F) and anti-Gβ (G) antibodies. GTPγS treatment activated Gα_o_-GTP as evidenced by its dissociation from Gβ.

### SDS-PAGE gel electrophoresis and Western blotting

For SDS-PAGE gel electrophoresis followed by total protein staining, IP samples were loaded on 4-12% Bis-Tris gels (NuPAGE #NP0322BOX) and separated using MOPS buffer (Novex #NP0001). Gels were stained with Imperial Protein Stain (Thermofisher Scientific #24615) per manufacturer’s instructions and images were captured using Epson Perfection V800 Photo Color Scanner. Images were processed by ImageJ software. For western blots, IP samples were loaded (20% of an IP sample/well) and separated on 10% SDS-PAGE gels. The protein was transferred onto a nitrocellulose membrane, and the blot was blocked and incubated with a primary anti-GOA-1 antibody (1:1000 diluted affinity-purified rabbit anti-GOA-1 polyclonal antibody (Patikoglou and Koelle 2002) at 4°C for overnight), washed, incubated with a secondary antibody (1:3000 HRP–linked Anti-Rabbit antibody Bio-Rad) and protein bands were visualized with SuperSignal West Pico PLUS Chemiluminescent Substrate (Thermofisher Scientific #34580) using a BioRad ChemiDoc MP system. Blots were reprobed for Gβ by stripping and then incubating overnight with 1:200 diluted mouse monoclonal anti-Gβ antibody (Santa Cruz #sc-166123) followed by a secondary incubation with 1:1000 diluted m-IgGk BP-HRP (Santa Cruz #sc-516102), and bands were again visualized by chemiluminescence.

### Reagent and data availability

Strains and plasmids used in this work are available upon request.

## Results

### Design of internally GFP-tagged Gα_o_ and expression in *C. elegans*

We designed a functionally-tagged *C. elegans* Gα_o_-GFP fusion protein, modeled in Figure 1A. The design inserts GFP that is flanked on either side by six amino acid flexible linkers into an internal loop of the alpha-helical domain of GOA-1, the *C. elegans* ortholog of Gα_o_ (Mendel et al., 1995; Ségalat et al., 1995). An analogous GFP insertion site was used by Hughes et al. (2001) to generate a mammalian Gα_q-_GFP fusion protein that was capable of mediating signaling in cultured cells when overexpressed with the α_2a_-adrenergic receptor. Gibson and Gilman (2006) further showed that insertion of YFP in the analogous loop of mammalian Gα_i1_ did not alter the nucleotide exchange or GTPase reaction rates of the purified protein, and that this Ga_i1_-YFP fusion protein could be activated in cultured cells by overexpressed and endogenous α_2_-adernergic receptors.

To express Gα_o_-GFP in *C. elegans*, we modified a 9.0 kb genomic clone containing the entire Gα_o_ gene (including its promoter, exons and introns, and 3’ region) to insert the GFP coding sequences between the codons for Gα_o_ amino acids T117 and E118. We also generated an activated mutant version of this Gα_o_-GFP clone in which we altered codon 205 to encode L instead of Q, a change that disrupts GTPase activity of Gα_o_ and renders the protein constitutively active (Mendel et al., 1995). The wild-type and Q205L versions of the Gα_o_-GFP transgene were separately inserted as single-copy transgenes into the *C. elegans* genome using Mos1 transposase (Frøkjaer-Jensen et al., 2008; Frøkjaer-Jensen et al., 2014). We also crossed the wild-type Gα_o_-GFP transgene into a mutant strain of C. elegans lacking endogenous Gα_o_ protein due to the Gα_o_ gene carrying an early stop codon mutation (Robatzek and Thomas, 2000).

Figure 1B shows Western blots of whole-worm protein lysates probed with an antibody against Gα_o_. The endogenous Gα_o_ protein and the higher molecular weight Gα_o_-GFP fusion proteins gave signals of similar intensity, suggesting that insertion of GFP into Gα_o_ did not interfere with its expression or stability.

We examined the localization of Gα_o_-GFP in *C. elegans* animals. Previous studies demonstrated that the *C. elegans* Gα_o_ gene is expressed in most or all neurons (Mendel et al., 1995; Ségalat et al., 1995). Using an antibody against *C. elegans* Gα_o_ to stain wild-type animals, we visualized endogenous Gα_o_ protein concentrated in bundles of neural processes, such as the nerve ring in the head, as well as in what appeared to be the plasma membranes of neural cell bodies (Figure 1C). Green fluorescence in transgenic animals carrying the Gα_o_-GFP or Gα_o_(Q205L)-GFP transgenes was localized in patterns that closely mimicked the localization of endogenous Gα_o_ (Figures 1D and 1E).

### Gα_o_-GFP rescues the locomotion and body morphology defects of a Gα_o_ null mutant

Gα_o_ null mutants have defects in locomotion behavior (Ségalat et al., 1995; Mendel et al., 1995), and we tested whether these defects could be rescued by transgenically-expressed Gα_o_-GFP. Figures 2A-2D show photographs of Petri plates on which individual worms have left tracks that reveal features of their locomotion behavior. We also used analyzed video of worms moving on such Petri plates to quantitate the abnormally deep body bends that Gα_o_ null mutants make during backwards locomotion (Figure 2E). Wild-type worms (Figure 2A) move forward with smooth sinusoidal body bends and rarely reverse direction, but Gα_o_ null mutants make abnormal body bends that leave abnormal tracks (Figure 2B) and that result in the animals sometimes bending so deeply during backwards locomotion that they touch their own body, events we term “reversal-touches” (Figure 2E). We found that expression of Gα_o_-GFP in the Gα_o_ null mutant background qualitatively rescued the Gα_o_ null mutant locomotion defects as judged by the tracks worms made (Figure 2C), and quantitation showed that the reversal-touch frequency defect was fully rescued (Figure 2D). The locomotion defects in Gα_o_ mutants arise at least in part because Gα_o_ is required to inhibit neurotransmitter release in ventral cord motor neurons that control locomotion behavior (Nurrish et al., 1999). The rescue of these defects suggests Gα_o_-GFP is functional in regulating neurotransmitter release.

Gα_o_(Q205L) is an activated mutant of Gα_o_ that is thought to block the GTPase activity of this G protein and thus traps Gα_o_ in its active GTP-bound state. Expression of Gα_o_(Q205L) in *C. elegans* leads to gain-of-function phenotypes, such as shallow body bends (Mendel et al., 1995). Transgenic expression of Gα_o_(Q205L)-GFP in otherwise wild-type *C. elegans* also caused a gain-of-function phenotype, since the tracks left by these worms show very shallow bends (Figure 2D), while similar transgenic expression of Gα_o_-GFP without the Q205L mutation did not have this effect (data not shown). Thus Gα_o_-GFP, like wild-type Gα_o_, can be activated by the Q205L mutation.

Gα_o_ null mutants have a scrawny body morphology which is seen in Figure 3A-3B and was also evident as a decrease in the length of the worms we photographed and tracked for the experiments shown in Figure 2A-2E. This body length defect was fully rescued by expression of Gα_o_-GFP (Figure 2F).

### Gα_o_-GFP partially rescues egg-laying behavior defects of a Gα_o_ null mutant

Gα_o_ null mutants show a hyperactive egg-laying behavior defect (Ségalat et a., 1995; Mendel et al., 1995; Koelle and Horvitz, 1996) due at least in part to Gα_o_ acting in the HSN motor neurons that stimulate egg laying to inhibit their release of serotonin (Tanis et al,. 2008). This defect leads Gα_o_ null mutant adult animals to retain very few unlaid eggs, since their eggs are laid almost as soon as they are produced (Figures 3A, 3B, and 3F). Another way to measure hyperactive egg-laying behavior is to count the fraction of freshly-laid eggs that are at early stages of development (Chase and Koelle, 2004). Hyperactive egg-laying mutants such as the Gα_o_ null mutant lay eggs so soon after they are fertilized that the laid eggs are often at the eight-cell stage or earlier, whereas wild-type animals rarely lay such early stage eggs (Figure 3G). Both assays of egg-laying behavior show that the hyperactive egg-laying defect in the Gα_o_ null mutant was substantially although not fully rescued by expression of Gα_o_-GFP (Figures 3C and 3F). The partial rescue of the Gα_o_ egg-laying defect seen in Figure 3 as opposed to the full rescue of the Gα_o_ locomotion reversal defect seen in Figure 2D may reflect differences in the ability of Gα_o_-GFP fully function in egg-laying versus locomotion neurons; alternatively, it could simply reflect an ability of the egg-laying assays to detect smaller differences in Gα_o_ function.

Transgenic expression of Gα_o_(Q205L)-GFP in otherwise wild-type *C. elegans* caused a gain-of-function phenotype in which animals fail to lay eggs, resulting in an accumulation of unlaid eggs, while similar transgenic expression of Gα_o_-GFP without the Q205L mutation did not have this effect (Figures 3A, 3D-3F). Thus Gα_o_-GFP not only rescues the behavioral defects of a Gα_o_ null mutant, but can also be activated by the Q205L mutation induce gain-of-function defects that are opposite to the defects of the null-mutant (Figures 2 and 3).

### Immunoprecipitation of Gα_o_-GFP in both its inactive and active states

Although an antibody that recognizes *C. elegans* Gα_o_ on Western blots (Patikoglou and Koelle, 2002) and in whole-mount stains of *C. elegans* animals (Figure 1C) is available, no antibody we have tested can be used to immunoprecipitate *C. elegans* Gα_o_, and this has remained an obstacle to biochemical studies of this protein. Therefore, we tested whether Gα_o_-GFP can be immunoprecipitated from *C. elegans* lysates using an anti-GFP monoclonal antibody. In these experiments, we assessed activation state of Gα_o_-GFP immunoprecipitated from whole-worm lysates by testing if the Gβ subunit co-precipitates, since Gβ should be in a complex with inactive but not with activated Gα_o_ (Figure 4A).

We found that Gα_o_-GFP, along with its associate Gβ subunit, were the major proteins found in anti-GFP immunoprecipitates of worm lysates expressing Gα_o_-GFP (Figure 4B-4D). When worm lysates expressing Gα_o_(Q205L)-GFP were immunoprecipitated with the same antibody, the Gα_o_-GFP protein but not Gβ were precipitated, confirming that the Q205L mutation locked Gα_o_-GFP in its active, GTP-bound state.

We were also able to fully activate the Gα_o_-GFP protein in a whole-worm lysate by incubating it with the non-hydrolysable GTP analog, GTPγS. The left lane of the gel in Figure 4E shows a total protein stain of a control immunoprecipitate of Gα_o_-GFP from a *C. elegans* lysate treated with GDP to maintain the G protein in an inactive state. An approximately equal amount of the Gα_o_-GFP and Gβ proteins were coprecipitated, as expected if the Gα_o_-GFP present in this lysate was close to 100% in the inactive, GDP-bound heterotrimer state in which Gα_o_-GFP and Gβ associate in a 1:1 stoichiometry, as modeled in Figure 4A. The identity of the major protein bands in Figure 4E as Gα_o_-GFP and Gβ were confirmed in the Western blots in Figures 4F and 4G. The right lanes in Figures 4E-4G show a parallel analysis of Gα_o_-GFP immunoprecipitates from the same protein lysate, but this time after treatment of the lysate with the non-hydrolysable GTP analog GTPγS. In this experiment, no detectable Gβ was coprecipitated with Gα_o_-GFP, consistent with the Gα_o_-GFP protein being close to 100% converted to the activated GTPγS-bound form, in which it is dissociated from Gβ as modeled in Figure 4A.

## Discussion

The goal of this work was to functionally tag Gα_o_ in *C. elegans* with a fluorescent protein to facilitate cell biological and biochemical studies of this major neural signaling protein. We adapted the approach of Hughes et al., (2001) by inserting GFP flanked by flexible linkers into a specific internal loop of the alpha-helical domain of the Gα protein. We found that single-copy transgenes express such Gα_o_-GFP fusion proteins in *C. elegans* at levels similar to that of the endogenous Gα_o_ protein, and that these Gα_o_-GFP fusion proteins appear to be localized to the processes and plasma membranes of neurons, similar to the localization of the endogenous Gα_o_ protein. The wild-type version of the Gα_o_-GFP protein can rescue the behavioral and body morphology defects of a Gα_o_ null mutant, although the extent of this rescue ranges from full to partial depending on the specific phenotypic defect analyzed. An activated mutant version, Gα_o_(Q205L)-GFP, can induce a gain-of-function phenotype. Immunoprecipitated wild-type Gα_o_-GFP appears to be stoichiometrically associated with Gβ, but can be fully activated and dissociated from Gβ either with the activating Q205L mutation or by incubation with the non-hydrolysable GTP analog GTPγS.

As of this writing, over 100 research articles have been published analyzing Gα_o_ function in *C. elegans*, reflecting the importance of this neural signaling protein (WormBase, http://www.wormbase.org, release WS279). Because this work has relied almost exclusively on genetic methods to make indirect inferences about the molecular mechanisms of Gα_o_ signaling, it has had a limited ability to make definitive conclusions about such mechanisms. For example, a several studies have speculated as to whether Gα_o_ signals by directly binding and activating the diacylglycerol kinase DGK-1 (the worm ortholog of mammalian DGKΘ based on genetic results consistent with this hypothesis (Nurrish et al., 1999; Miller et al., 1999; Jose and Koelle, 2005; Koelle, 2018); however, the lack of methods to immunoprecipitate Gα_o_ protein complexes from worm lysates have prevented a clear test of this hypothesis. Another line of genetic work in *C. elegans* has suggested that major neural protein kinase CaMKII (known as UNC-43 in *C. elegans*) may phosphorylate Gα_o_ to regulate Gα_o_ signaling (Robatzek and Thomas, 2000), but this hypothesis has not been tested using biochemical approaches for lack of tools to isolate and directly examine the *C. elegans* Gα_o_ protein from lysates of wild-type versus *unc-43* mutants.

Beyond its role in neural signaling, Gα_o_ also plays a central role in mitotic spindle positioning during asymmetric cell divisions in early development (Gotta and Ahringer, 2001). Cell biological studies of asymmetric cell division in *C. elegans* embryos have depended heavily on the use of functional GFP fusions to proteins that control cell polarity and mitotic spindle positions, as these tools make it possible to track the positioning and dynamic movements of these proteins during cell divisions (Rose and Gönczy, 2014). Antibody stains suggest that Gα_o_ may be localized to spindle asters in dividing embryonic cells (Gotta and Ahringer, 2001), but this early finding has not been followed up. Our development of functional Gα_o_-GFP transgenes should enable a more definitive analysis of the role of Gα_o_ in asymmetric cell divisions.

## Acknowledgements

This work was supported by National Institutes of Health Grants NS036918 and NS086932 to M.R.K. Strains were provided by the *Caenorhabditis* Genetics Center, funded by the National Institutes of Health Office of Research Infrastructure Programs P40 OD010440. We thank Nakeirah Christie and Halie Sonnenschein for editing the manuscript.

